# Suppression of ERK signalling promotes pluripotent epiblast in the human blastocyst

**DOI:** 10.1101/2024.02.01.578414

**Authors:** Claire S. Simon, Afshan McCarthy, Laura Woods, Desislava Staneva, Qiulin Huang, Madeleine Linneberg-Agerholm, Alex Faulkner, Athanasios Papathanasiou, Kay Elder, Phil Snell, Leila Christie, Patricia Garcia, Valerie Shaikly, Mohamed Taranissi, Meenakshi Choudhary, Mary Herbert, Joshua M. Brickman, Kathy K. Niakan

## Abstract

Studies in the mouse demonstrate the importance of fibroblast growth factor (FGF) and extra-cellular receptor tyrosine kinase (ERK) in specification of embryo-fated epiblast and yolk-sac-fated hypoblast cells from uncommitted inner cell mass (ICM) cells prior to implantation. Molecular mechanisms regulating specification of early lineages in human development are comparatively unclear. Here we show that exogenous FGF stimulation leads to expanded hypoblast molecular marker expression, at the expense of the epiblast. Conversely, we show that specifically inhibiting ERK activity leads to expansion of epiblast cells functionally capable of giving rise to naïve human pluripotent stem cells. Single-cell transcriptomic analysis indicates that these epiblast cells downregulate FGF signalling and upregulate molecular markers associated with naïve pluripotency. Our functional study demonstrates for the first time the molecular mechanisms governing ICM specification in human development, whereby segregation of the epiblast and hypoblast lineages occurs during maturation of the mammalian embryo in an ERK signal-dependent manner.

## Introduction

Formation of the blastocyst is a critical event in embryogenesis occurring in the first week of human development. Two sequential cell fate decisions segregate the embryonic pluripotent epiblast from the extra-embryonic tissues, trophectoderm and hypoblast. Errors in cell differentiation can cause embryo arrest, leading to miscarriage. More than half of all natural conceptions are estimated to end in very early (<5 week) pregnancy loss^1^. Despite the significance for human health and stem cell biology, we do not understand the mechanisms which direct cell differentiation in the early human embryo. To improve our understanding of the mechanisms of cell fate specification in the early embryo, we studied the effects of cell-cell communication during this critical window of development.

The ability of FGFs to bias ICM cells towards hypoblast is conserved amongst mammals, including in rodents, rabbits, bovine and pig^2–5^. The molecular pathway governing ICM cell fate segregation has been elucidated genetically in the mouse. Within mouse ICM cells, and later in epiblast cells, the pluripotency marker NANOG induces *Fgf4* expression^6^. FGF4 ligand then signals in an autocrine manner to uncommitted ICM progenitors predominantly via FGFR1, then later by FGFR2 in hypoblast precursors^7, 8^. Signalling activation then specifies hypoblast via a GRB2/MEK/ERK cascade leading to an upregulation of hypoblast specific markers such as *Gata6*, and a downregulation of epiblast markers like *Nanog*^7–11^. These molecular insights have also informed strategies to establish naïve mouse pluripotent stem cells *in vitro* that more closely resemble the blastocysts stage epiblast *in vivo* from which they were derived^12, 13^. However, in human embryos, previous studies suggested that upstream FGF receptor (FGFR) or mitogen-activated protein kinase (MEK) inhibition did not affect hypoblast formation^3, 14^.

We recently demonstrated crosstalk of FGF-driven mitogen-activated protein kinase (MEK) and insulin growth factor-driven phosphoinositol-3 kinase (PI3K) activity upstream of ERK signalling in human pluripotent stem cells (hPSCs) derived from the epiblast^15^. In addition, we showed that hypoblast differentiation *in vitro* from naïve hPSCs depends on FGF signalling and that naïve hPSCs can be maintained by simultaneously blocking the FGF receptor and its downstream kinase MEK^16^. We hypothesized that ERK may have a conserved role in human embryo hypoblast versus epiblast specification.

Here, we determined that exogenous FGF signalling activity is sufficient to specify the hypoblast in human blastocysts. We describe ERK signalling during the blastocyst stage and demonstrate that blocking ERK activity leads to expansion of epiblast cells functionally capable of giving rise to naïve hPSCs. Transcriptomic analysis further reveals that these epiblast cells downregulate FGF signalling, while maintaining molecular markers of the epiblast. Our functional studies provide mechanistic insight into human blastocyst formation and reveal for the first time the molecular mechanisms that regulate ICM specification in humans. We propose a unified model in which segregation of the epiblast and hypoblast lineages occurs during maturation of the mammalian blastocyst in an ERK signal-dependent manner.

## Results

### Exogenous FGF is sufficient to drive human hypoblast specification

We hypothesized that FGF4 may play a conserved role in human preimplantation development. Transcriptional analysis of human pre-implantation embryos^17–19^ indicates that there is pan-ICM expression of the FGFR1 receptor, while FGF4 is expressed specifically in epiblast cells (Supplementary Fig. 1a), consistent with current mouse models of ICM segregation where FGF4 acts predominantly via FGFR1 upstream of pERK in uncommitted ICM cells to specify hypoblast^7, 8^.

To test the effect of exogenous FGF signalling on human embryos we carried out a dose-response experiment. We treated Day 5 human embryos and cultured them for 36h in medium containing 0ng/ml, 250ng/ml, 500ng/ml, 750ng/ml and 1000ng/ml FGF4 plus Heparin (which stabilizes FGF interactions) (Fig. 1a). Immunofluorescence analysis of NANOG (epiblast), GATA4 (hypoblast) and GATA3 (trophectoderm, placental progenitor cells) showed that increasing concentrations of FGF4 increased the number of hypoblast cells (Figs. 1a,1b and Supplementary Fig. 1b). The mean number of hypoblast cells in 750ng/ml FGF4 treated embryos increased by 2-fold, when compared to control embryos (p = 0.04, mean 16 vs 8 hypoblast cells per embryo) (Fig. 1b). While there was a 2-fold reduction in the mean number of epiblast cells when comparing 750ng/ml FGF4 treated embryos with controls, this difference was not significant (Fig. 1c, p = 0.07, mean 3 vs 7 epiblast cells per embryo). Increasing concentrations of FGF4 had no significant effect on ICM cell number (Supplementary Fig. 1c). Consistent with changes in both epiblast and hypoblast numbers, we found that increasing concentration of FGF4 increased the ratio of hypoblast : epiblast cells in the ICM (Fig. 1d, Control 46% hypoblast, 750ng/ml 84% hypoblast p=0.0004 and 1000ng/ml 84% hypoblast p=0.01), and embryos exhibited an all-hypoblast ICM in a dose-dependent manner (Supplementary Fig. 1d).

**Fig. 1.**
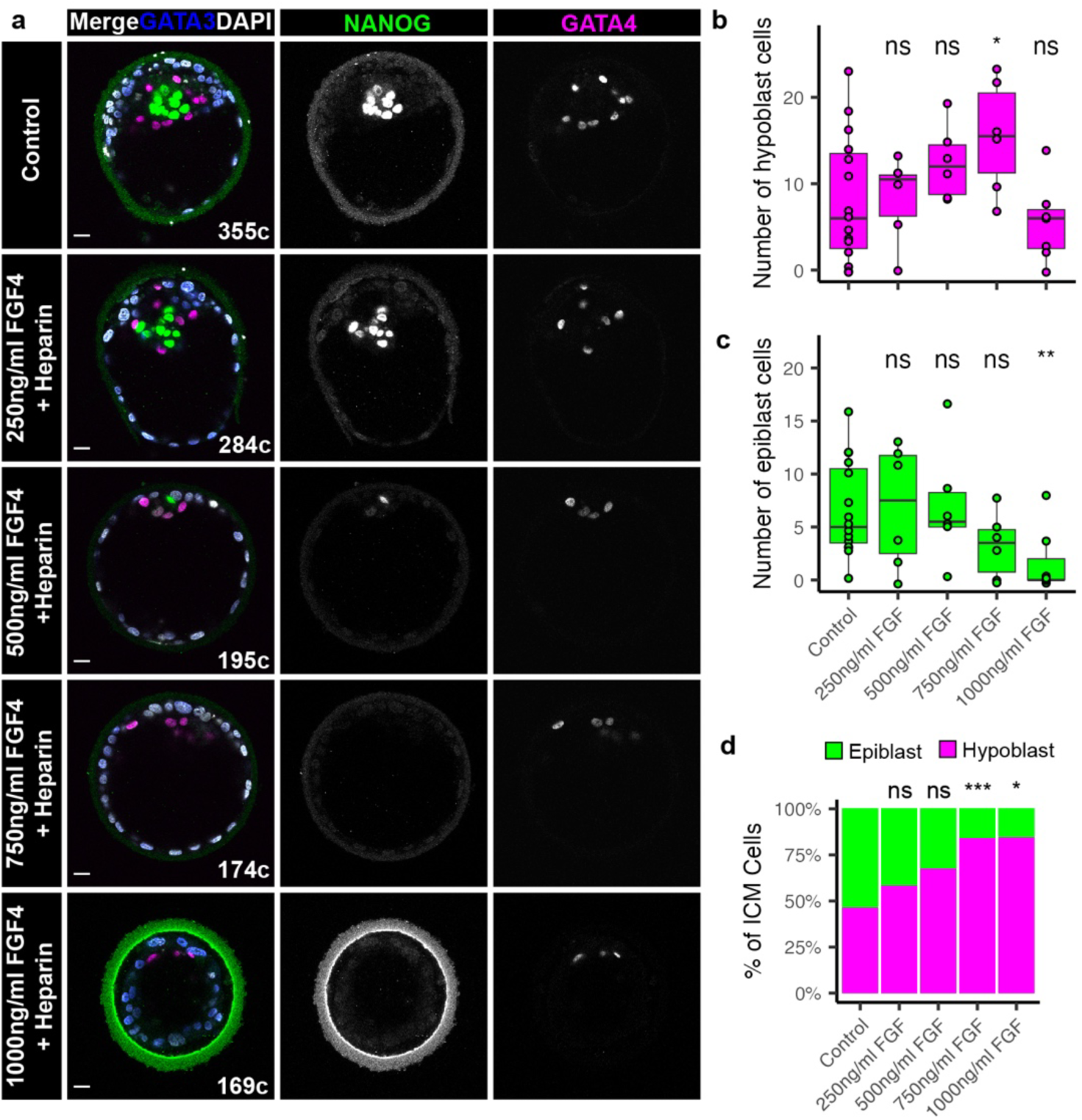
Exogenous FGF is sufficient for driving human hypoblast specification. **(a)** Confocal images of Day 6.5 human embryos immunofluorescently labelled for lineage markers NANOG (epiblast), GATA4 (hypoblast) and GATA3 (trophectoderm), and stained for nuclear DAPI. Human embryos were cultured in Control medium, or medium supplemented with increasing concentrations of FGF and Heparin as indicated from Day 5 for 36 hours. Total cell number (c) indicated. Scale bars 20 µm. **(b - c)** Boxplots showing the number of **(b)** hypoblast (GATA4+NANOG-GATA3-) and **(c)** epiblast (NANOG+GATA4-GATA3-) cells in human embryos cultured in increasing concentrations of FGF and Heparin. Values for each embryo are shown as individual points. Control n = 15, 250ng/ml n = 7, 500ng/ml n = 6, 750ng/ml n=6, 1000ng/ml n=7. **(d)** Stacked bar charts showing the mean proportion of epiblast and hypoblast in the ICMs per embryo in each treatment group. Embryos without ICMs were excluded from the analysis. Control n = 14, 250ng/ml n = 5, 500ng/ml n = 6, 750ng/ml n=6, 1000ng/ml n=6. Two-tailed t-test, n.s. not significant * p < 0.05, ** p < 0.01, *** p < 0.001.

This is consistent with the response in mouse and cow embryos to similar doses of FGF4, where NANOG or SOX2 (epiblast) expression is downregulated and the ICM is predominantly comprised of SOX17- or GATA6-expressing hypoblast cells (Supplementary Fig. 1e, 1f), similar to previous studies^2, 3^. Moreover, we determined that rat preimplantation embryos also responded to FGF4, downregulating NANOG and upregulating GATA6 expression throughout the ICM (Supplementary Fig. 1g). Overall, our findings demonstrate that FGF4 is sufficient to drive human hypoblast specification, a mechanism conserved across species.

### Suppression of ERK signalling blocks hypoblast formation in the human blastocyst

FGFs can activate multiple downstream pathways, including ERK, PKC and PI3K ^20^. Our previous work identified active phosphorylated ERK (pERK) in human whole blastocyst protein lysates^15^. However, we had not defined the cell type that contains this signalling activity. To address this question, we analyzed pERK by immunofluorescence staining^21^, beginning with mouse embryos to establish a positive control. Consistent with previous studies^21, 22^, pERK protein expression is detectable in the ICM (Supplementary Fig. 2a). We next characterized pERK expression by immunofluorescence analysis in primed human embryonic stem cells (hESCs) and detected pERK cytoplasmic localization. As expected, MEK inhibition using PD0325901 led to downregulation of pERK in hESCs, indicating specificity of the immunofluorescence analysis (Supplementary Fig. 2b). Next, we stained for pERK in human blastocysts from Days 5 to 6.5, using SOX2 and OTX2 as markers of the epiblast and hypoblast, respectively (Fig. 2a). Active pERK is evident in the cytoplasm of epiblast, hypoblast and trophectoderm cell types, with a high proportion of hypoblast having ERK^high^ cells (Fig. 2b). The level of pERK in embryos increases over time (Fig. 2c), altogether suggesting a role for this signalling pathway in early human development.

**Fig. 2.**
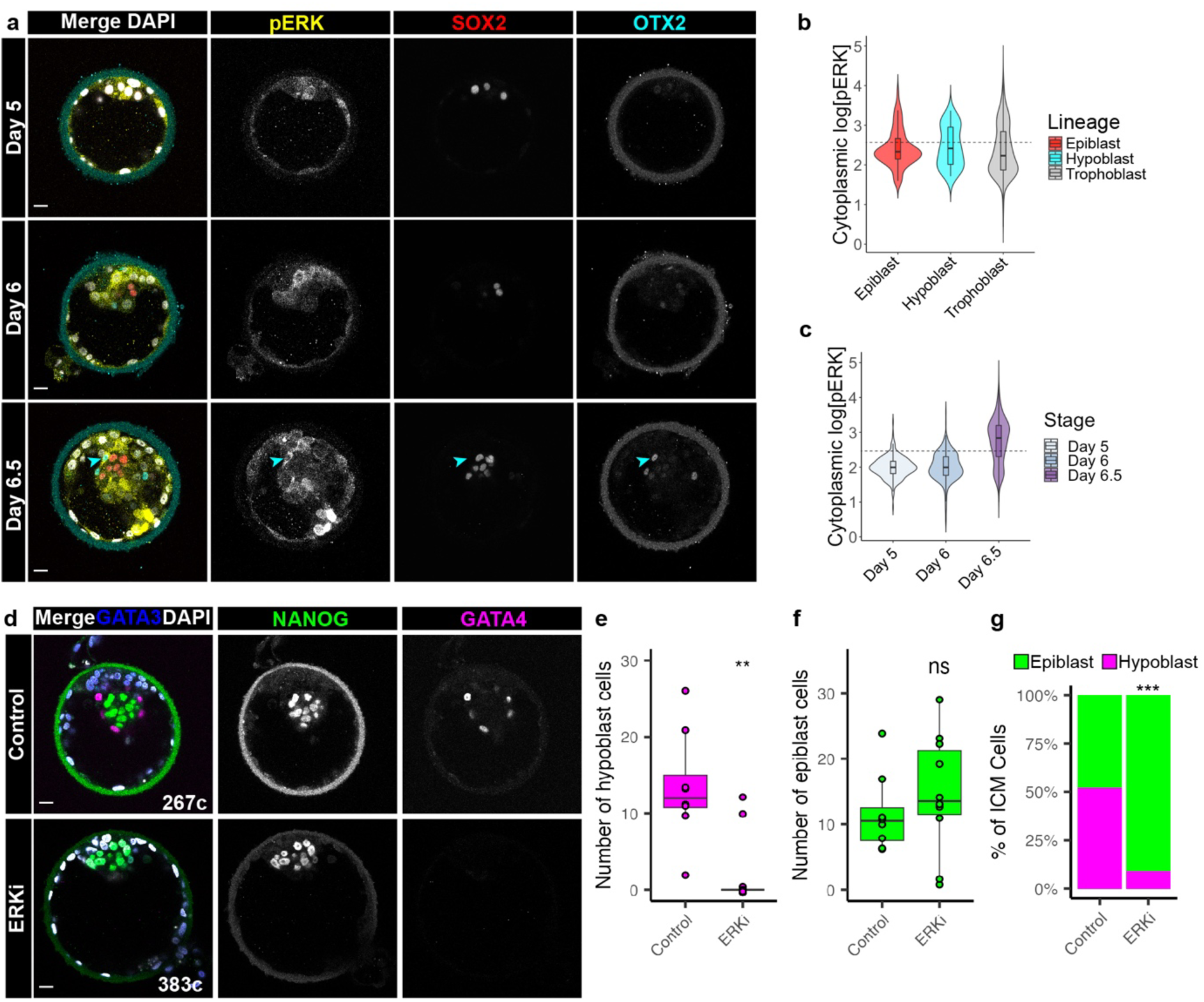
Suppression of ERK signalling blocks hypoblast formation in the human blastocyst. **(a)** Confocal images of Day 5 - 6.5 human embryos immunofluorescently labelled for phophorylated (p)-ERK, lineage markers SOX2 (epiblast), OTX2 (hypoblast), and stained for nuclear DAPI. Cyan arrow indicates a hypoblast cell with high pERK levels. Scale bars 20 µm. **(b)** Violin plots showing the cytoplasmic fluorescence intensity of pERK in each lineage in embryos shown in (a). Epiblast n = 83 cells, hypoblast n = 41 cells, trophectoderm n = 1535 cells; n = 10 embryos. Dashed line indicates threshold between ERK^low^ and ERK^high^ cells. **(c)** Violin plots showing the cytoplasmic fluorescence intensity of pERK over time in embryos shown in (a). Day 5 n = 313 cells; 3 embryos, Day 6 n = 542 cells; 3 embryos, Day 6.5 n = 804 cells; 4 embryos. Dashed line indicates threshold between ERK^low^ and ERK^high^ cells. **(d)** Confocal images of Day 6.5 human embryos immunofluorescently labelled for lineage markers NANOG (epiblast), GATA4 (hypoblast) and GATA3 (trophectoderm) and stained for nuclear DAPI. Human embryos cultured with or without ERKi (5µm Ulixertinib) from Day 5 for 36 hours. Total cell number (c) indicated. Scale bars 20 µm. **(e-f)** Boxplots showing the number of (e) hypboblast (GATA4+) and (f) epiblast (NANOG+) cells in human embryos cultured with and without ERKi. Values for each embryo are shown as individual points. **(g)** Stacked bar charts showing the mean proportion of epiblast and hypoblast in the ICMs per embryos in each treatment group. Control n = 8, ERKi n = 10. Two-tailed t-test, ns= not significant, * p < 0.05, ** p < 0.01, *** p < 0.001

To determine if ERK is the effector of ICM specification, we used Ulixertinib ^23^, a selective ATP-competitive inhibitor of ERK1/2. As expected, Ulixertinib treatment (ERK inhibition treatment, referred to as ERKi) led to downregulation of GATA4 hypoblast in the mouse and an ICM exclusively comprised of NANOG expressing epiblast cells (Supplementary Data Fig 2c), consistent with previous studies^24^. In addition, the ICM of cow embryos following ERKi lost SOX2-GATA6+ hypoblast cells (Supplementary Fig. 2d).

We then cultured Day 5 human embryos for 36h in medium containing volume matched DMSO (Control) or 5µM Ulixertinib (ERKi) (Fig. 2d) followed by quantitative immunofluorescence staining (Supplementary Fig. 2e). Blocking ERK signalling resulted in embryos with no GATA4 expression, and an ICM comprising of predominantly NANOG positive cells (Figs. 2d, 2e, 2f). ERKi treatment thus led to a loss of hypoblast in the majority of treated embryos (Fig. 2e, p = 0.003, 2 vs 13 hypoblast cells per embryo). Additionally, there was a modest increase in the number of epiblast cells in ERKi embryos compared with controls (Fig. 2f, p = 0.4, 15 vs 12 epiblast cells per embryo). The proportion of hypoblast : epiblast cells was dramatically distorted in ERKi treated embryos as there were no significant changes in ICM cell numbers (Fig. 2g, Supplementary Fig. 2f, 2g, control 52% hypoblast, and ERKi 9% hypoblast, p= 0.0002). The majority of ERKi embryos demonstrated an all-epiblast ICM phenotype (Supplementary Fig. 2g), while those that retained some hypoblast cells also had a greater cell number indicating a more advanced developmental age, and suggesting this residual hypoblast could be a result of lineage specification that occurred prior to exposure to ERKi.

We also observed a similar phenotype when culturing embryos for a shorter window of 24 hours, from Day 5 in ERKi. Here, ERKi embryos lost the early hypoblast marker GATA6 and retained high levels of pluripotency epiblast markers NANOG and OCT4 (Supplementary Fig. 2h). Thus, human ICM specification to epiblast and hypoblast likely occurs between the early- (Day 5) to mid- (Day 6) blastocyst stages. Together, these results show ERK signalling is active at the time of ICM segregation in human blastocysts, and suppression of this pathway blocks hypoblast formation.

### ERKi of human embryos leads to upregulation of transcripts associated with naïve pluripotency

To characterize the impact of ERK signalling inhibition on epiblast versus hypoblast lineage segregation, we performed single-cell RNA-seq analysis on ERKi treated human embryos compared to DMSO treated controls. We integrated this dataset with reference untreated control blastocyst datasets at embryonic days 6 and 7^17–19^, which are equivalent stages of development to the treatment conditions.

To investigate transcriptional differences between the reference and control versus ERKi-treated cells, we focused on cluster 2, which is enriched for ICM cells that express epiblast and hypoblast molecular markers^25^ (Figs. 3a, 3b, 3c, Supplementary Data 3a). Differential gene expression analysis of cells in ICM cluster 2 indicated that ligands and receptors of the FGF and downstream ERK signalling pathway (e.g. *FGF4*, *GRB2*, *FGFR3*, *FGF23*), ERK-induced signal regulated ETS transcription factors (*ERF*, *ETV1*, *ETV4*, *ETV5*, *ETV6*), and downstream regulators (e.g. *DUSP6*, *DUSP7*, *DUSP8*, *DUSP9*, *DUSP14*, *SPRY4*) were transcriptionally affected following ERKi (Fig. 3d, Supplementary Table 1). Moreover, ERKi-treated ICM cells downregulated *GATA6*, *GATA4*, *PDGFRA*, *HNF4A*, *SOX17*, *DAB2*, *COL4A1*, and immediate early FGF responses genes, such as *DUSP6*, indicating that both signalling and hypoblast-enriched transcripts were perturbed (Fig. 3d, Supplementary Table 1). By contrast, transcripts associated with the pluripotent epiblast or naïve pluripotency such as *DNMT3L*, *DNMT3A*, *DPPA5*, *KLF17*, *KLF3*, *KLF5*, *NODAL*, *POU5F1*, *PRDM1*, *PRDM14* and *SOX2*, were significantly upregulated in ICM cells following ERKi-treatment compared to controls (Fig. 3d, Supplementary Table 1).

**Fig. 3.**
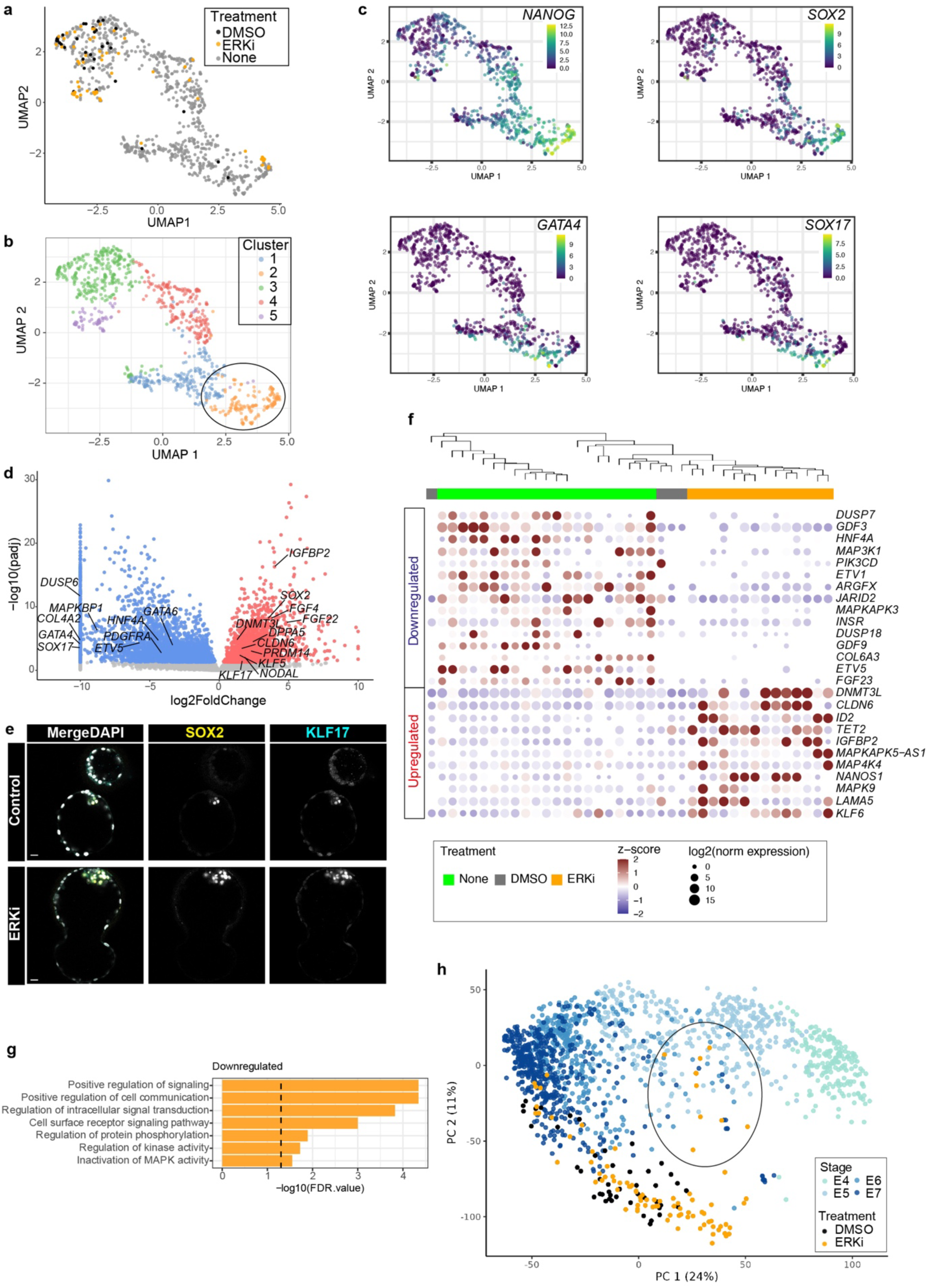
ERKi of human embryos leads to upregulation of transcripts associated with naïve pluripotency. **(a)** UMAP of embryonic cells at days 6 - 7, calculated based on lineage markers from a previous publication^25^. No treatment reference control data^17–19^ (grey), human embryos cultured in ERKi (orange) or volume-matched DMSO (black). **(b)** UMAP coloured according to cluster identity. **(c)** UMAP coloured by expression of epiblast or hypoblast marker genes from previous publication^25^. Scales show log transformed normalised expression values. **(d)** Volcano plot with genes significantly differentially expressed (DESeq2 padj < 0.05) between ERKi treated and control (DMSO and untreated) cells from the ICM cluster 2 in panel A. Representative genes upregulated (red) or downregulated (blue) DESeq2 padj < 0.05 in ERKi compared to control (DMSO and untreated) cells are highlighted. X axis shows the mean log2 fold change (ERKi – control) with large values capped at 10. **(e)** Confocal images of Day 6.5 human embryos immunofluorescently labelled for SOX2 and KLF17 (epiblast) and stained for nuclear DAPI. Scale bars 20 µm. **(f)** Hierarchical clustering of DMSO and ERKi treated cells and selected high-confidence epiblast control samples from previous publication^26^. Selected downregulated or upregulated genes (DESeq2 padj < 0.1) are shown and labeled as log2 normalized and z-score expression values. **(g)** Representative gene set enrichment analysis of terms associated with genes that were significantly downregulated in ERKi versus control cells in the epiblast cluster. **(h)** Principal component analysis of single cells from embryonic days 4 - 7 from previous studies^17–19^ (shades of blue), DMSO treated controls (black) and ERKi-treated (orange). Principal components calculated based on the top 10% variable genes (n = 957).

We confirmed at the protein level that the pluripotent epiblast markers SOX2 and KLF17 are expressed throughout the ICM and that the early hypoblast marker PDGFRA is downregulated following ERKi-treatment (Fig. 3e, Supplementary Fig. 3b). This is consistent with our previous results where NANOG and OCT4 pluripotency markers are similarly retained throughout the ICM in ERKi-treated embryos (Fig. 2d, Supplementary Fig. 2h).

To investigate transcriptional differences specifically between the ERKi-treated versus control epiblast cells within the ICM, we transcriptionally compared the cells we sequenced in this study to a recently curated high-confidence set of epiblast, hypoblast and trophectoderm cells^26^ (Supplementary Fig. 3c, Supplementary Table 2). This indicated that ERKi and DMSO treated cells were transcriptionally similar to either epiblast or trophectoderm and that we did not collect any hypoblast cells in either treatment condition (Supplementary Fig. 3c). Differential gene expression analysis comparing epiblast reference and control versus ERKi-treated epiblast-like cells confirmed that components of the FGF/ERK signalling pathway were significantly disrupted. This included downregulation of FGF ligands (*FGF5* and *FGF23*), the downstream MAP kinase *MAPKAPK3*, negative feedback regulators (*DUSP6* and *DUSP18*), and transcriptional effector ETS-family member, *ETV5* (Fig. 3f, Supplementary Fig. 3d, Supplementary Table 2). Pathway enrichment analysis further shows a significant effect on genes associated with signal transduction, including inactivation of MAPK activity (Fig. 3g). Notably, genes associated with Insulin/PI3K signalling were also both transcriptionally up and downregulated (e.g. *INSR, IGF2, IGFBP2. MAP4K4, MAPK9* and *FOS*) (Fig. 3f, Supplementary Fig. 3d, Supplementary Table 2).

We noted that while molecular markers associated with naïve pluripotency were upregulated, markers of formative/primed pluripotency such as *ETV5*, *FGF5* and *WNT7B* were significantly downregulated in ERKi-treated cells compared to controls (Supplementary Fig. 3d, Supplementary Table 2). We speculate that abolishing FGF/ERK activity may lead to a more immature epiblast state in ERKi cells, similar to observations in mouse^7, 8, 12^. We therefore integrated single cell datasets from embryonic days 4 and 5 of development, representing the morula and early blastocyst stages of development, respectively (Fig. 3h). Principal component analysis indicated that 9 out of the 14 ERKi treated epiblast cells were transcriptionally more similar to early blastocyst cells at embryonic day 5 (Fig. 3h, Supplementary Fig. 3e, 3f). Altogether this indicates that ERKi treatment leads to transcriptional downregulation of ERK signalling components and upregulated naïve pluripotency associated molecular markers.

### Suppression of ERK signalling promotes pluripotent epiblast identity

To functionally test the pluripotent potential of the epiblast following ERKi treatment of embryos, we aimed to derive naïve hESC lines. Previously, naïve hESCs, the *in vitro* stem cell counterpart of the pre-implantation epiblast, were derived directly from isolated ICM cells in conditions containing a titrated concentration of GSK3, MEK and PKC inhibitors together with LIF (t2iLGö) medium ^27–29^. We initially tested to see if we could directly derive naïve hESCs from human blastocysts in the further optimized medium where the GSK3 inhibitor has been replaced with XAV939, a tankyrase inhibitor and Wnt pathway antagonist (PXGL)^30^. We plated Day 6 human blastocysts, intact or following mural trophectoderm laser dissection. We were able to derive 5 stable naïve hESC cell lines from 8 blastocysts in both intact and dissected conditions (Supplementary Data Figs. 4a-c), demonstrating efficient naïve hESC derivation in PXGL medium.

After 36h in either ERKi or DMSO culture, we plated whole intact blastocysts in naïve PXGL medium to derive naïve hESCs (Supplementary Fig. 4d). After ICM outgrowth and passaging, we were able to establish 2/8 naïve hESC lines from control embryos and 5/9 naïve hESC lines from the ERKi treated embryos (Fig. 4a). These lines had characteristic compact and domed naïve ESC morphology (Fig. 4b), and immunofluorescence analysis showed they expressed core pluripotency markers OCT4 and SOX2, and naïve pluripotency markers KLF4, KLF17, SUSD2, and TFAP2C^31–34^ (Figs. 4c, 4d).

**Fig. 4.**
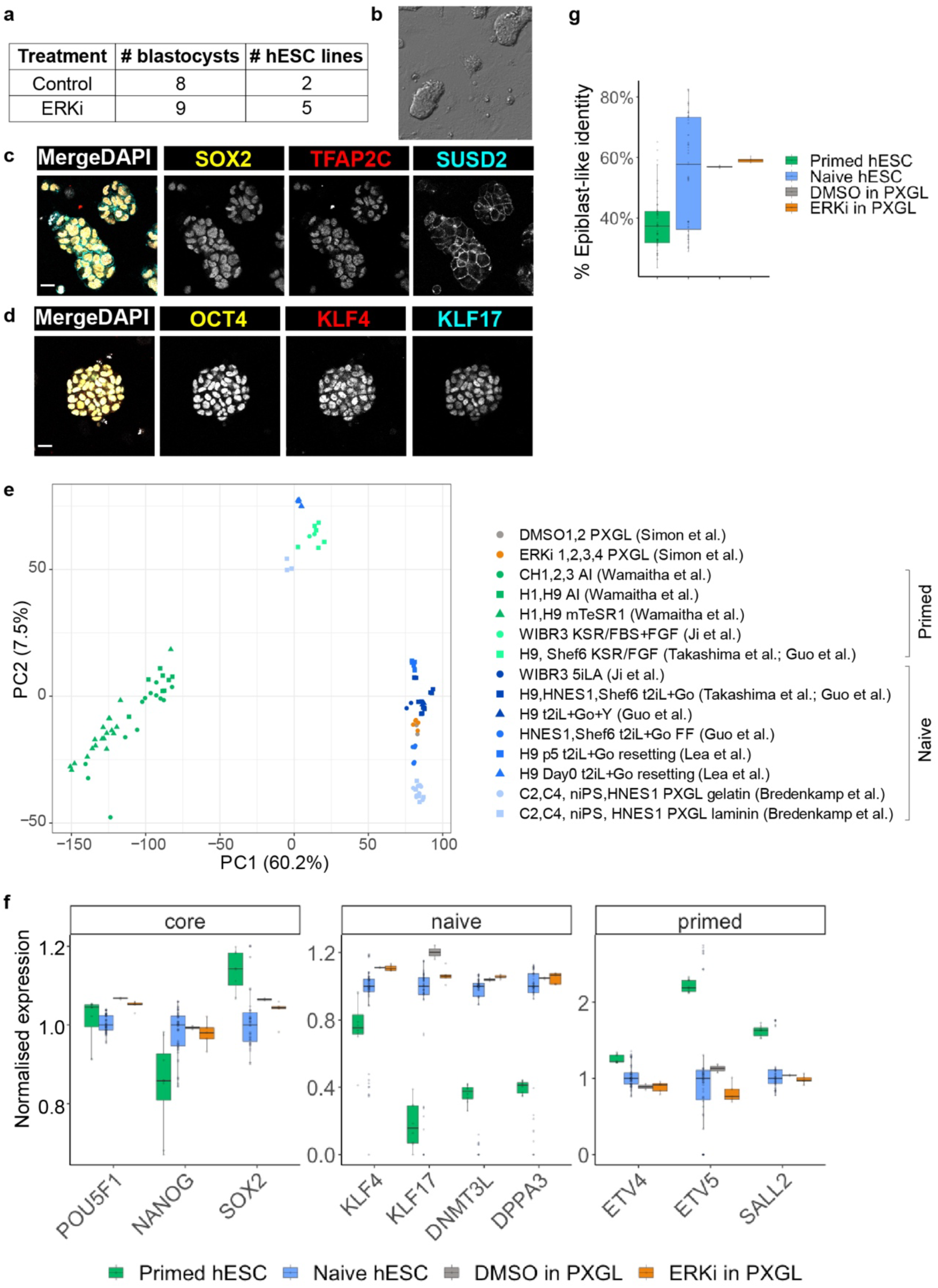
Suppression of ERK signalling promotes pluripotent epiblast identity. **(a)** Table showing outcome of naïve hESC derivation from Control and ERKi cultured embryos. **(b)** Phase-contrast image of a naïve hESC line derived from an ERKi embryo. **(c-d)** Confocal images of naïve hESC line derived from an ERKi embryo, immunofluorescently labelled for naïve (TFAP2C, SUSD2, KL4, KLF17) and core (SOX2, OCT4) pluripotency markers and stained for nuclear DAPI. Scale bars 20µm. **(e)** Principal component analysis following RNA-seq of hESC. The reference cell lines used are noted in the legend compared to the PXGL hESC lines derived in this study. Per-gene variance was modeled, and significantly variable genes (DESeq2 padj < 0.05) were used in the loading. **(f)** Boxplot showing the normalized expression (Log2 of the DESeq2 normalized read counts) of selected pluripotency genes relative to median expression within published naïve PSCs. Previously published primed (n = 48) or naïve (n = 55) hESCs or in the PXGL hESC lines (n = 2 of DMSO PXGL hESCs or n = 5 for ERKi PXGL) derived in this study. **(g)** Boxplot of genes enriched in epiblast cells were used as a comparison to previously published primed (n = 48) or naïve (n = 55) hESCs or PXGL hESC lines derived in this study following ERKi (n = 5) or DMSO (n = 2) treatment.

The hESCs derived in PXGL media following either ERKi or DMSO treatment were transcriptionally more similar to previously established naïve hPSCs than primed hPSCs (Fig. 4e). Moreover, the PXGL hPSCs were similar to previously derived naïve hPSCs in their high expression of naïve markers *KLF4*, *KLF17*, *DNMT3L*, and *DPPA3*, low expression of primed markers *ETV4, ETV5* and *SALL2* and equivalent level of general pluripotency markers *POU5F1*, *NANOG* and *SOX2* compared to primed hPSCs (Fig. 4f). PXGL hPSCs derived following ERKi or DMSO treatment also resembled previously established naïve hESCs in transcriptional similarity to human blastocyst epiblast cells compared to primed hPSCs (Fig. 4g). Furthermore, the newly established PXGL hPSCs cell lines were karyotypically normal (Supplementary Fig. 4e).

We next sought to determine if the MEK inhibitor PD0325901 (“P” in PXGL media) in naïve pluripotent stem cell derivation and maintenance could be replaced with our ERKi Ulixertinib (“U” in UXGL media). Following 36h of either ERKi or DMSO treatment we plated the laser dissected ICM of blastocysts in UXGL media (Supplementary Fig. 4f). Following ICM outgrowth and subsequent passaging, we were able to establish 1/4 naïve hESC from control embryos and 2/4 naïve hESC lines from ERKi treated embryos (Supplementary Fig. 4g). As above, the UXGL hESC lines had characteristic domed shaped morphology and expressed expressed SOX2, NANOG and OCT4 in addition to naïve pluripotency markers KLF4, KLF5, KLF17, TFAP2C and DNMT3L (Supplementary Data Figs. 4h, 4i). In conclusion, ERKi embryos retain pluripotency markers throughout their ICM and give rise to ground-state pluripotent stem cells, demonstrating their naïve potential.

## Discussion

Our data demonstrates that FGF4 is sufficient to drive human hypoblast specification in a dose dependent manner. Divergence of epiblast and hypoblast markers between FGF4 expressing and non-expressing cells in the human ICM further supports FGF/ERK governing lineage segregation^35^. However, it is still an outstanding question which ligands drive this signalling activity *in vivo*. Further work is needed *in vivo* to confirm endogenous human FGF4 function in hypoblast specification, and whether other RTKs such as IGFs^15^, or parallel pathways such as WNTs as has been suggest in marmoset monkeys^14^, may additionally be involved.

The necessity for FGF/ERK in human hypoblast specification has also been suggested by stem cell models, where conversion of naïve ESC to hypoblast-like naïve endoderm (nEnd) is prevented by FGFR inhibition^16^. However, attempts at blocking the FGF/MEK/ERK pathway using small molecule inhibitors against MEK or FGFR produced varying outcomes in mammalian embryos, and might argue for intrinsic species-specific differences in the molecular mechanism of ICM cell fate decisions. For example, while mouse, rat and rabbit embryos all lost hypoblast cells upon MEKi, only mouse and rat have an accompanying expansion of epiblast cells^2, 5, 14^. By contrast, in pig, cow, and marmoset embryos MEKi did not completely block hypoblast specification, but the numbers of cells were reduced^3, 4, 36^. Most surprisingly, in human embryos, MEKi and/or FGFRi did not affect hypoblast formation in a significant manner^3, 14^. One possibility for the discrepancy is that the concentrations of the FGFR- or MEK-inhibitors used, ratio of media to mineral oil or diffusion across the zona may all impact on efficacy of the treatment and were therefore insufficient to fully inhibit ERK activity^37^. Moreover, we speculate that there may be compensation or crosstalk from other pathways^15^ whereby FGFR or MEK inhibition may not have completely abolished ERK activity. A recent study using human stem cell-based embryo models, and exogenous FGF2 or FGFR inhibition treatment of human embryos, corroborates the findings we present here, suggesting a role for FGF in hypoblast specification^38^.

We find that inhibiting ERK signalling leads not only to expansion of epiblast marker expression, but also upregulation of genes associated with naïve pluripotency competent for naïve hESCs derivation. Moreover, we replaced the requirement for MEKi with an ERKi, thereby establishing an alternative method for naïve hPSC derivation. It will be interesting to determine in the future if these cells are an improvement over previously established naïve hPSCs in terms of karyotypic stability, capacitation, directed differentiation or epigenetic characteristics such as genomic imprinting.

Our comparison across mouse, rat, cow and human embryos demonstrates a conserved mechanism regulating the second cell fate specification decision shortly after fertilization. Our data further supports step-wise specification of three lineages in human preimplantation development. This is consistent with our functional data revealing the importance of cell polarity and Hippo signalling activity in regulating ICM versus trophectoderm cell fate specification^39, 40^. Our data is also consistent with recent re-analysis of single-cell RNA-seq data together with protein analysis characterizing ICM progenitor cells, suggesting a step wise progression of TE versus ICM specification followed by segregation of the ICM to epiblast and hypoblast cells^41, 42^. Altogether we show that ERK activity can alter the balance between EPI and hypoblast, suggesting a cell fate switch of a common ICM progenitor. These insights inform our understanding of human development and stem cell biology.

## Materials and Methods

### Ethics statement

This study was approved by the UK Human Fertilisation and Embryology Authority (HFEA): research licence numbers R0162, R0397, R0401 and R0152 and independently reviewed by the Health Research Authority’s Research Ethics Committee IRAS projects 308099, 252286 and 272218.

The process of licence approval entailed independent peer review along with consideration by the HFEA Licence and Executive Committees and the Research Ethics Committee. Our research is compliant with the HFEA Code of Practice and has undergone multiple inspections by the HFEA since the licence was granted.

Informed consent was obtained from all couples that donated spare embryos following infertility treatment. Before giving consent, people donating embryos were provided with all the necessary information about the research project, an opportunity to receive counselling and the conditions that apply within the licence and the HFEA Code of Practice. Donors were informed that embryos used in the experiments would be stopped before 14 days post-fertilization and that subsequent biochemical and genetic studies would be performed. Informed consent was also obtained from donors for all the results of these studies to be published in scientific journals. No financial inducements were offered for donation. Consent was obtained for creation and culture of embryonic stem cell lines from these embryos and deposition of cell lines in the UK Stem Cell Bank. Embryos surplus to the patient’s IVF treatment were donated cryopreserved and were transferred to the University of Cambridge and Francis Crick Institute, where they were thawed and used in the research project.

### Human Embryo Thaw

Human embryos were thawed using vit kit-Thaw (Fujifilm) for vitrified embryos, or BlastThaw^TM^ Kit (CooperSurgical) for slow frozen embryos according to the manufacturer’s instructions.

### Human Embryo Culture

Human blastocysts were cultured in pre-equilibrated Global Media (Cooper Surgical) supplemented with 10% Human Serum Albumin (HSA, Cooper Surgical) in Embryo+ slide dishes (Vitrolife) overlaid with mineral oil (Cooper Surgical) at a ratio of 1 media : 9 mineral oil. Embryos were alternatively cultured in GT-L culture media. Embryos were incubated at 37°C 5.5% CO_2_ in an EmbryoScope time-lapse imaging system (Vitrolife). For treatments, immediately after thawing embryos were incubated in Global media, 10% HSA supplemented with; 250 FGF: 250ng/ml FGF4 (R&D) and 250ng/ml Heparin (Sigma); 500 FGF: 500ng/ml FGF4 and 500ng/ml Heparin; 750 FGF: 750ng/ml FGF4 and 750ng/ml Heparin; 1000 FGF: 1000ng/ml FGF4 and 1000ng/ml Heparin; 5µM Ulixertinib (Cambridge Bioscience); 1% DMSO. Day 5 embryos were treated with cytokines and inhibitors for either 36 or 24 hours.

### Human and Mouse Embryo Dissociation

Human and mouse embryos were subject to immunosurgery to remove outer trophectoderm cells and isolate the inner cell mass (ICM), as reported previously^27, 43^. The zona pellucida was removed by washing embryos through drops of Acidic Tyrode’s solution (Merck) pre-warmed to 37°C and overlaid with mineral oil. Embryos were then washed briefly and incubated at 37°C for 30 min in 20% anti-mouse serum antibody (Sigma) or 20% anti-human serum antibody (Sigma), depending on the species, in Global media + 10% HSA. Embryos were then washed briefly in Global Media + 10% HSA, before incubation with 20% Guinea Pig Serum Complement (Merck) at 37°C for 10-15 min until trophectoderm cell began to lyse. Embryos were then moved to a drop of medium and triturated with 100 µm STRIPPER tip (Cooper Surgical) to remove lysed trophectoderm cells and to isolate the inner cell mass.

Isolated ICMs were then briefly washed, then incubated at 37°C for 3 min in 0.5% Trypsin, 1 mM EDTA in PBS (Fisher Scientific). Then, ICMs were transferred to 4% BSA, 0.5 mM EDTA in PBS for fine manual dissociation into single cells with pulled (Sutter Instruments) glass capillaries (World Precision Instruments). Cells were then picked manually with pulled glass capillaries for downstream single-cell RNA-seq.

### Single cell RNA-seq

Single-cell cDNA synthesis was performed using SMART-Seq v4 Ultra Low Input RNA Kit for Sequencing (Takara) according to the manufacturer’s protocol with some modifications as previously described^17^. Briefly, single-cells were manually picked after dissociation and snap frozen on dry ice in 5 µl 10x Reaction Buffer in low-bind 0.2 ml PCR tubes and stored at −80°C until processing. Samples were subject to first strand cDNA synthesis, then cDNA was amplified by LD-PCR for 23 cycles. Amplified cDNA was purified on AMPure XP beads, using a 96-well magnetic stand (ThermoFisher), and eluted in 17 µl of elution buffer. Single-cell RNA-seq libraries were prepared using a Nextera XT DNA Library Preparation Kit (Illumina) according to the manufacturer’s protocol. Sequencing was carried out as paired-end 50 bp read on Novaseq 6000 to a depth of approximately 12 million reads per cell.

### scRNA-seq processing and analysis

Single-cell RNA-seq data was pre-processed as described in Alanis-Lobato et al 2023^26^, https://github.com/galanisl/early_hs_embryo_GRNs. We integrated scRNA-seq data from 3 different preimplantation human embryo studies profiling the three cell types present at the late blastocyst stage^17–19^, with the DMSO and ERKi treated samples generated in this study. High confidence lineage annotations for the cells in the previous studies were obtained from previous work^25^.

Technical replicate fastq files were merged and pre-processed using the nf-core/rnaseq pipeline 3.10.1 with Nextflow 22.10.1^44^. Briefly, reads were trimmed for adaptors and quality using Trimgalore 0.6.7 and cutadapt 3.4. Trimmed reads were pseudoaligned to GRCh38 with salmon 1.9.0 and used to generate estimated raw abundances and TPM normalized reads. Resulting gene expression matrices were integrated, normalized, and log2 transformed using Bioconductor tools^45^. Datasets were filtered to retain high-quality cells with > 20000 reads, > 200 genes detected, and < 90% of reads mapping to mitochondrial genes. Mitochondrial genes and genes with an average count < 1 across all samples were removed prior to normalization.

To analyse lineage specification of E6-E7 cells; gene-level filtering, normalization, scaling, and log-transformation were repeated on a restricted set of cells at stages E6 and E7. Twelve marker genes representing the epiblast, primitive endoderm, and trophoblast lineages were used for PCA^25^. The resulting first 5 principal components were used as the input for UMAP analysis, with the number of neighbors set to 50. Clusters were detected based on the PCA output, using clusterCells^46^ with 25 nearest neighbors.

Epiblast cells were selected based on the clustering output and tested for differential expression between control (DMSO treated samples from this study, and untreated samples from previous publications^17–19^, versus Ulixertinib treatment.

Differential expression between the ERKi and control samples was determine using DESeq2 with the following options: test=“LRT”, useT=T, minmu=1e-6, minReplicatesForReplace=ing, fitType=“glmGamPoi”.

To analyse the developmental trajectory of E4-E7 cells, per-gene variance was modelled with scran^46^ and the top 10% most variable genes (n = 957) were used for principal component analysis (PCA).

### Human primed ESC culture

Primed H9 human ESC were cultured according in mTeSR1 (StemCell Technologies) according to the manufacturer’s guidance, on Matrigel (356231, BD Bioscience) coated plates at 37°C 5% CO_2_ under normoxia. H9 hESCs were and dissociated using ReLeSR (Stem Cell Technologies) incubate for 4 min at 37°C and passaged in clumps.

### Human naïve ESC culture

Human naïve cells were cultured in PXGL medium^30^: N2B27 medium (Takara, Y40002), supplemented with 1uM PD0325901 (Cambridge Bioscience; 13034-1mg-CAY), 2uM XAV939 (Cambridge bioscience, CAY13596-1mg), 2 µM Gö6983 (Bio-Techne; 2285/1), 10ng/ml hLIF (PeproTech; 300-05), and Pen/Strep (Gibco), on MEF feeder layer (∼1 x 10^5^ cells / cm^2^) and incubated under hypoxic conditions 5%O_2_ 5%CO_2_ 37°C in a humid incubator. Culture medium was replaced every other day, and cells were passaged every 4-5 days with a split ratio of 1:3 – 1:5. Cells were passaged by washing with PBS, followed by incubation with Accutase (Thermo, A1110501) at 37°C for 5-10 min with gentle pipetting and monitoring to confirm single-cell dissociations. Cells were washed and pelleted, before resuspension and culture in PXGL+ROCKi medium for 24 h on a fresh MEF feeder plate. After 24 h, the medium was replaced with fresh PXGL (without ROCKi). Alternatively, the 1 µM PD0325901 above was replaced with 5 µM Ulixertinib (Cambridge Bioscience) to generate UXGL medium.

### Human naïve ESC derivation

Human Day 5 embryos were cultured for 36 hours in ERKi or in DMSO control medium or thawed on Day 6 and cultured for 2 hours in Global medium. Embryos that had not yet hatched had the zona pellucida removed by a brief wash in Acidic Tyrode’s pre-warmed at 37°C. Whole intact embryos, or laser dissected embryos to remove mural trophectoderm, were then plated on MEF coated 4-well plates (Nunc) in pre-equilibrated PXGL or UXGL medium. PXGL or UXGL medium was made from N2B27 medium (Takara, Y40002), supplemented with either 1 µM PD0325901 (Cambridge Bioscience; 13034-1mg-CAY) or 5 µM Ulixertinib (Cambridge Bioscience) plus: 2 µM XAV939 (Cambridge bioscience, CAY13596-1mg), 2 µM Gö6983 (Bio-Techne; 2285/1), 10 ng/ml hLIF (PeproTech; 300-05), and Pen/Strep (Gibco). Embryo outgrowths were left undisturbed for 48 hours at 5%O_2_ 5%CO_2_ 37°C, then fed with pre-equilibrated half medium changes every 2 days. After 7-10 days, outgrowths were manually picked under a dissecting microscope. Outgrowths were dissociated by washing, and then incubated in microdrops of Accutase under mineral-oil for 5 min at 37°C before trituration into single cells with 290 µm and 100 µm STRIPPER tips (CooperSurgical). Dissociated single cells were then washed in PXGL or UXGL medium and plated onto MEF feeder plates with PXGL or UXGL medium supplemented with ROCK inhibitor 10 µM Y27632 (Tocris), and then re-fed with PXGL or UXGL medium every 2 days. Clones were passaged manually every 5-7 days until sufficient numbers for bulk passaging, initially in a 1:1 split ratio, then 1:2 – 1:3 split ratio every 3-5 days.

### Bulk RNA-seq

Stable naïve hESC lines were used for the bulk RNA-seq experiments. Human naïve ESCs were dissociated to single cells. 6×10^5^ cells were resuspended in 600 µl RLT buffer (Qiagen) and snap frozen on dry ice for later processing. RNA was isolated using the RNAeasy Mini Kit (Qiagen, 74104), QIAShredder DNase digest (Qiagen, 79654), automated on a QIAcube (Qiagen) according to the manufacturer’s instructions. The RNA library prep was performed using the mRNA polyA (NEB) kit, before paired-end 100 bp sequencing on a NovaSeq2 (depth of 25M reads).

### hESC RNA-seq analysis

Reads were assessed for quality with FastQC and trimmed with TrimGalore 0.5.0. Reads were pseudoaligned to GRCh38 using Salmon 0.11.3. Mitochondrial genes, pseudogenes, ribosomal genes, and genes not detected in any condition were removed. Samples were normalised using DESeq2 and invariant genes were removed. Any single-cells present were subject to imputation using DRImpute. Per-gene variance was modelled with scran 1.26.2 and variable genes FDR < 0.05 were used as input into principal component analysis. Similarity to an epiblast identity was assessed using DeconRNASeq^47^, using as a reference set high confidence epiblast cells that were obtained from previous work^26^.

### Karyotyping

To determine chromosome copy number, 2 x 10^5^ cells from each of the hESC lines derived in this study were collected. To extract DNA the DNeasy® Blood & Tissue Kit was used according the manufacture’s protocol (Qiagen, 69504 and 69506). This was followed by low-pass next generation sequencing (depth of sequencing < 0.1x). Libraries were prepared using the VeriSeq PGS Kit (Illumina) or the NEB Ultra II FS Kit according to the manufacturer’s instructions. The MiSeq platform or the Illumina HiSeq 4000 platforms were used. Reads were aligned to the human genome hg19 using BWA v0.7.17^48^ and the copy number profiles generated with QDNaseq v1.24.0 as described previously^49^.

### Mouse embryo culture

All animal research was performed in accordance with the UK Home Office regulations under project licence PP8826065, which passed ethical review by the Francis Crick Institute Animal Welfare Review Board in 2019. Three- to -four to eight-week-old (C57BL6 × CBA) F1 female mice were super-ovulated using injection of 5 IU of pregnant mare serum gonadotrophin (PMSG; Sigma-Aldrich). Forty-eight hours after PMSG injection, 5 IU of human chorionic gonadotrophin (HCG; Sigma-Aldrich) was administered. Superovulated females were set up for mating with eight-week-old or older (C57BL6 × CBA) F1 males. Mouse zygotes were isolated in FHM under mineral oil and cumulus cells were removed with hyaluronidase (Sigma-Aldrich; H4272). Mouse embryos were cultured in pre-equilibrated Global Media (Cooper Surgical) supplemented with 10% Human Serum Albumin (HSA, Cooper Surgical) overlaid with mineral oil (Cooper Surgical) at a ratio of 1 media : 9 mineral oil.

### Human embryo immunofluorescence staining

Human embryos were washed briefly in PBS-/- + 1% HSA, followed by fixation in 4% PFA (diluted from 16% paraformaldehyde solution (Fisher Scientific) on ice for 1 h. Embryos, were then washed three times in PBS-/- + 0.1% Triton X-100 (PBX; Sigma), and permeabilized in PBS-/- + 0.5% TritonX-100 for 20 min. Embryos were washed briefly in PBX, then incubated in blocking solution: 10% Donkey serum (Jackson ImmunoResearch, 017-000-121) in PBX, for 1 hour on rotating shaker. Embryos were incubated with primary antibodies (Supplementary Table 3) diluted in blocking solution overnight at 4°C on rotating shaker. The next day, embryos were washed three time in PBX, and incubated in blocking solution for 1 hour on rotating shaker, before incubating with secondary antibodies diluted in blocking solution for 1 hour on rotating shaker. Finally, embryos were washed four times in PBX, and a final wash PBS-/- + 0.1% Triton + 3.33% Vectashield with DAPI (Vector Laboratories, H-1200). For staining of pERK, embryos were stained as above, with the following modifications: 1 x PhosSTOP (Roche, 4906845001) was included in all solutions up to and including the primary antibody incubation, embryos were fixed in 8% PFA + 1x PhosSTOP at room temperature for 10 min on rotating shaker. Primary and secondary antibodies used in this study are outlined in Supplementary Table 3.

### Bovine IVF and culture

Oocytes were collected from bovine ovaries and matured overnight in BO-IVM medium (IVF Bioscience) at 38.5°C under normoxia. Frozen bull sperm straws (UK Sire Services) were thawed for 1min at 37°C, then transferred to BO-SEMENPREP (IVF Bioscience) followed by two rounds of washes and centrifugation at 300g for 5min. Granulosa cells were removed from cumulus-oocyte complexes (COCs) by pipetting and combined with 4×10^6^ spermatozoa per 1ml of BO-IVF media (IVF Bioscience) for 6-10hrs at 38.5°C under normoxia. COCs were then denuded and transferred to BO-IVC media (IVF Bioscience) overlayed with mineral oil at 20-30 presumptive embryos per 500µL drop followed by culture at 38.5°C under hypoxia. At 7dpf, the embryos were transferred to BO-IVC media supplemented with either 5µM Ulixertinib or 750ng/mL FGF4 and 1µg/mL heparin and cultured for an additional 48hrs.

### Immunofluorescence of bovine embryos

Bovine blastocysts were fixed with 100mM Hepes pH 7, 50mM EGTA pH 7, 10mM MgSO_4_, 4% methanol-free formaldehyde and 0.2% Triton X-100 in sterile H_2_O for 30min at 37°C followed by 3x washes with PBS with 3mg/mL polyvinylpyrrolidone (PVP). Fixed blastocysts were permeabilised with 0.5% Triton X-100 in PBS with 3mg/mL PVP and blocked with 3% donkey serum, 0.1% bovine serum albumin and 0.2% Triton X-100 in PBS with 3mg/mL PVP, both overnight at 4°C. Primary antibodies were diluted in blocking buffer and incubated overnight at 4°C and secondary antibodies were diluted 1:500 in blocking buffer and incubated for 2hrs at room temperature shaking. The embryos were washed 3x for 10min following each antibody incubation. DAPI was added to blocking buffer during the wash steps after secondary antibody incubation for nuclear stain. Bovine blastocysts were then imaged in blocking buffer without DAPI in 18-well ibidi slides using a Leica SP8 confocal microscope.

### Human embryo fluorescence imaging

Fixed samples were imaged on a Leica SP8 scanning confocal microscope or a Zeiss LSM880. Embryos were mounted in microdroplets of PBS + 0.025% Tween20 + 1.5% Vectashield, on glass bottomed 35mm dishes (Maktek, P35G-1.5-14-C) coated with mineral oil. Embryos were imaged along the entire z-axis with a 1µm step using a glycerol immersion HC PLAN APO CS2 63x 1.30 NA objective (Leica) or a 40x or 20x objective (Zeiss). Imaging parameters were kept consistent across experiments.

### Human ESC immunofluorescence

Primed and naïve human ESC were washed briefly in PBS, before fixation in 4% PFA at room temperature for 10 minutes. Cells were washed three times in PBS + 0.1%Triton-X (PBX) before 20 min permeabilization in 0.5% Triton X-100 on a rotating shaker. Cells were then washed, incubated in blocking solution (10% Donkey serum in PBX) for 1 hr at room temperature. Then, cells were incubated overnight in primary antibody in blocking solution at 4 degrees. Cells were washed three times in PBX, before a second incubation in block solution for 1hr, followed by incubation with secondary antibodies. Antibodies outlined in Supplementary Table 3.

### Image processing and quantification

Raw images were processed in ImageJ. Nuclear segmentation for quantification of nuclear and cytoplasmic fluorescence was carried out using Stardist and CellProfiler as described previously^32^. Correction of segmentation errors and classification of inner ICM vs outer TE cells was performed manually. Adjustments for fluorescence decay along the z-axis was performed by linear regression followed by empirical Bayes method, as detailed in Saiz et al^50, 51^.

## Acknowledgments

We thank the donors of human embryos whose contributions were essential for this research. We thank all members of the Niakan lab for their technical assistance and for help and comments on the manuscripts. We thank the Centre for Trophoblast Research for technical support and advice. We thank Florian Holfelder and Timo Kohler for support with confocal microscopy. We thank the Cambridge Stem Cell Institute for access to cell culture reagents.

We thank the Francis Crick Institute’s Science Technology Platforms: Lyn Healy and Liani Devito from the Human Embryo and Stem Cell Unit; Advanced Sequencing Facility, Advanced Light Microscopy and the Genomics Equipment Park. Work in the laboratory of KKN was supported by the Wellcome 221856/Z/20/Z (KKN). Work in the laboratories of KKN and MH was supported by the Wellcome Human Developmental Biology Initiative 215116/Z/18/Z. Work in the laboratory of KKN was also supported by the Francis Crick Institute which receives its core funding from Cancer Research UK FC001120, the Medical Research Council FC001120 and Wellcome FC001120. For the purpose of Open Access, the authors have applied a CC BY public copyright licence to any Author Accepted Manuscript version arising from this submission.

## Author contributions

Conceptualization: CSS; Methodology: CSS, AM, KKN; Investigation: CSS, AM, AF, KKN, DS, QH; Visualisation: CSS, KKN; Analysis: CSS, LW, AF; Funding acquisition: KKN, MH; Project administration: CSS, KKN; Supervision: CSS, KKN; Human embryo consenting and facilitating donations: TP, KE, PS, LC, SA, VS, MT, MH, MC; Writing – original draft: CSS, KKN; Writing – review & editing: CSS, KKN with support from all authors.

## Competing interests

Authors declare that they have no competing interests.

## Data and materials availability

Raw data produced for this study has been uploaded onto GEO accession GSE250614: https://www.ncbi.nlm.nih.gov/geo/query/acc.cgi?acc=GSE250614 A reviewer token is needed for access. Details of the analysis methods and code are available at: https://gitlab.developers.cam.ac.uk/ctr/ctr-bioinformatics/niakan-lab/kkn21_cs564_001. Single-cell RNA-seq data from E4-E7 embryos were downloaded from GEO accessions GSE66507 and GSE36552, and ArrayExpress https://www.ebi.ac.uk/biostudies/arrayexpress/ accession E-MTAB-3929. Primed and naive hESC datasets were downloaded from the ENA Browser https://www.ebi.ac.uk/ena/browser/home accessions PRJEB7132, PRJNA286204, PRJNA522065, PRJNA575370, PRJEB12748, and PRJEB47485. All reagents, codes and materials used will be available to any researcher for the purposes of reproducing or extending the analysis.

## Notes

### Competing Interest Statement

The authors have declared no competing interest.

